# Choice-Induced Preference Change under the Value Refinement Framework

**DOI:** 10.1101/2022.07.15.500254

**Authors:** Douglas G. Lee, Giovanni Pezzulo

## Abstract

Sequential sampling models of choice, such as the drift-diffusion model (DDM), are frequently fit to empirical data to account for a variety of effects related to accuracy/consistency, response time (RT), and sometimes confidence. However, no model in this class can account for the phenomenon known as *choice-induced preference change*, where decision makers tend to rate options higher after they choose them and lower after they reject them. Studies have reported choice-induced preference change for many decades, and the principal findings are robust. The resulting *spreading of alternatives* (SoA) in terms of their subjective value ratings is incompatible with traditional sequential sampling models, which consider the rated values of the options to be stationary throughout choice deliberation. Here, we propose an extension of the basic DDM that allows the drift rate to vary across deliberation time depending on which attributes are attended to at which points in time. Critically, the model assumes that choice deliberation commences based only on the more salient attributes of the options, and that additional attributes eventually come into consideration when the decision cannot be resolved based on the initial attributes alone. We show that this model can account for SoA (in addition to choice consistency and RT), as well as all previously reported relationships between SoA and choice difficulty, attribute disparity, and RT.

## Introduction

A considerable proportion of contemporary research on value-based decision-making focuses on choices between two options, using subjective ratings of the individual options (obtained in a separate task) as independent variables to help explain choice behavior. This research has exposed many robust patterns with respect to both choice response (i.e., which option the agent chose from the set of available options) and response time (RT; the temporal delay between the agent becoming aware of the choice options and declaring its response). First among such patterns is that so-called difficult decisions (those between options of similar subjective value) are generally more stochastic (i.e., predictions about which option will be chosen are less accurate) and take longer to conclude, relative to so-called easy decisions (Milosavljevic et al., 2010; Ratcliff & McKoon, 2008). Decisions are also thought to be more difficult when the value estimates of the choice options are less certain, and such decisions result in qualitatively similar behavioral patterns as those deemed difficult based on value difference (Gwinn & Krajbich, 2020; Lee & Coricelli, 2020; Lee & Daunizeau, 2020, 2021; Lee & Hare, 2023). Another common pattern is that decisions for which the value of the choice set as a whole is larger are usually made more quickly, and often more consistently (Hunt et al., 2012; Polanía et al., 2014; Sepulveda et al., 2020; Shevlin et al., 2022; Shevlin & Krajbich, 2021; S. M. Smith & Krajbich, 2019; Teodorescu et al., 2016). Finally, choices between options with high attribute disparity (e.g., where each option dominates within a different attribute dimension) are associated with faster RT (Bhatia & Mullett, 2018; Lee & Hare, 2023; Lee & Holyoak, 2021). Each of these independent variables (absolute value difference, value certainty, value sum, and attribute disparity) also has a robust positive impact on choice confidence (the subjective belief an agent has about whether it chose the best option from the choice set; (Brus et al., 2021; Folke et al., 2016; Lee & Hare, 2023)).

Beyond choice consistency, RT, and confidence, there is another variable commonly measured in experimental choice paradigms. Though largely overlooked in the decision-making literature, the phenomenon known as *choice-induced preference change* has been well documented in the social psychology literature (Bem, 1967; Brehm, 1956; Festinger, 1957; Harmon-Jones & Mills, 2019) and more recently demonstrated in value-based decision-making studies (Alós-Ferrer et al., 2012; Chammat et al., 2017; Colosio et al., 2017; Coppin et al., 2010, 2014; Egan et al., 2007, 2010; Greenberg & Spiller, 2016; Hagège et al., 2018; Ito et al., 2019; Izuma et al., 2010, 2015; Johansson et al., 2014; Koster et al., 2015; Lieberman et al., 2001; Luo & Yu, 2017; Miyagi et al., 2017; Nakamura & Kawabata, 2013; Salti et al., 2014; Sharot et al., 2009, 2010, 2012; Shultz et al., 1999; Tandetnik et al., 2021; Taya et al., 2014; Voigt et al., 2017, 2019); see (Enisman et al., 2021) for a meta-analysis). With so-called choice-induced preference change, a preference (direction and/or magnitude) between options that is implicitly inferred via isolated value ratings differs before versus after the preference that is explicitly observed via choices between pairs of options. The key variable here is often referred to as the *spreading of alternatives* (SoA), because it typically entails the value estimates of the choice options spreading apart as the decision maker reassesses them during or after deliberation (see also the literature on *coherence shifts* and *information distortion*: (Carlson & Russo, 2001; Holyoak & Simon, 1999; Russo et al., 1996, 2008; D. Simon et al., 2001, 2004, 2008)). Studies such as those listed above have shown that SoA is a robust phenomenon and is particularly high when choices are difficult (e.g., the difference in option values and/or the certainty about option values is low) or when attribute disparity is high (Lee & Hare, 2023; Lee & Holyoak, 2021). It has also been shown that SoA and RT are strongly negatively correlated, and that SoA and choice confidence are strongly positively correlated (Lee & Coricelli, 2020; Lee & Daunizeau, 2020, 2021; Lee & Hare, 2023; Lee & Holyoak, 2021, 2023).

### Sequential sampling model accounts of choice behavior

Various computational models of simple decisions (e.g., two-alternative forced-choice) exist, the most common of which come from the sequential sampling / accumulation-to-bound class (Brown & Heathcote, 2008; Busemeyer & Townsend, 1993; Ratcliff et al., 2016; Usher & McClelland, 2001). Under this type of model, the agent processes information about the choice options incrementally until the so-called evidence in favor of one of the options passes a threshold and that option is declared to be the winner (and is therefore chosen). Although multiple variations of this core model class have been proposed, they generally rely on a relative-value accumulator that tallies information about the options based on random samples from underlying probability distributions (one for each option) until it reaches a preset evidence threshold / response boundary. This class of model has provided an elegant account of choice and RT patterns in a variety of domains, in particular with respect to the impact of choice difficulty (value proximity; see (Lee & Usher, 2023) for a version that also includes value uncertainty as a component of choice difficulty). With some minor modifications, this type of model has also simultaneously accounted for the impact of overall set value on choice behavior (Krajbich et al., 2010; Lee & Usher, 2023; Shevlin et al., 2022; Shevlin & Krajbich, 2021; S. M. Smith & Krajbich, 2019). Even choice confidence can be accounted for by certain variations of sequential sampling model, either as a secondary readout of the evidence accumulation process (Calder-Travis et al., 2021; De Martino et al., 2013; Kiani et al., 2014; Moran et al., 2015; Moreno-Bote, 2010; Pleskac & Busemeyer, 2010; Ratcliff & Starns, 2009; van den Berg et al., 2016; Vickers & Packer, 1982; Zylberberg et al., 2012) or as the primary readout itself (Lee, Daunizeau, et al., 2023).

The evidence accumulation-to-bound framework has proven to be a powerful tool, yet there remains an important gap in its explanatory power. Despite the success that it has had in accounting for most relevant variables in the study of simple decisions (i.e., accuracy/consistency, RT, and sometimes confidence), it remains fundamentally incapable of accounting for the spreading of alternatives phenomenon, by virtue of the very assumptions that it is built on. In brief, sequential sampling models rely on stationary inputs to the evidence accumulation process, typically regarded to be the means of latent probability distributions representing the value of each option in the mind of the decision maker. The fact that these inputs are assumed to be stationary precludes the possibility that such models could account for changes in value estimates.

### Sequential sampling models and the spreading of alternatives

Recent interest has arisen in finding a way to address the spreading of alternatives (SoA) with sequential sampling models (Lee & Daunizeau, 2021; Zylberberg et al., 2024). Lee and Daunizeau suggest that SoA occurs as a result of a tradeoff between mental effort and choice confidence (Lee & Daunizeau, 2020, 2021). When faced with a difficult decision (e.g., if the options seem equally valuable at first and thus initial confidence about knowing the best option is low), a decision maker might prefer to invest mental effort towards processing additional information with the intention to distinguish the option values more confidently, rather than making an immediate choice with low confidence. The cognitive processes that eventually result in SoA would therefore be instrumental to the decision as it unfolds since they would effectively increase the discriminability of the options and make the choice easier. To address SoA, this proposal removes a key assumption of classical sequential sampling models — namely, that preference responses develop based on value-related signals that are *retrieved* (e.g., from memory) and compared rather than being *constructed* during deliberation (see the literature on *constructed preferences*: (Ariely et al., 2006; DeKay et al., 2011; Lichtenstein & Slovic, 2006; Payne et al., 1999; D. Simon et al., 2008; Warren et al., 2011)).

The model of Tajima and colleagues (Tajima et al., 2016), which contends that evidence accumulation summarizes a Bayesian updating of option values, could offer a different explanation of SoA (as prior estimates evolve to posterior estimates). This model assumes that both options are initialized with null (or at least equal, if there is prior knowledge about the summary statistics of the population of potential options) prior value estimates, but that the value estimates can come to differ as they are elucidated over time. From this perspective, preferences are not constructed during deliberation and do not change over time; they are simply more clearly recognized as their signals stabilize. This could result in an observed SoA effect between the start and end of deliberation. However, empirical SoA is calculated using subjective value ratings obtained both prior to and after the actual choice. It is unclear how the Tajima et al model might account for ratings that were systematically different when elicited after versus before the choice. Furthermore, this model would hold that value estimates constructed during choice deliberation could never shift closer together from start to end of deliberation, since they are initialized identically. Empirically, value estimates often do shift closer together, as evidenced by negative SoA on individual trials (Lee & Holyoak, 2021, 2023). Importantly, this would also mean that decision makers could never change their minds between start and end of deliberation, because all potential preferences would be null at decision onset.^1^ Previous studies have shown such changes of mind (i.e., choices in favor of options that were initially rated lower, but later rated higher than the alternatives) to be common (Lee & Daunizeau, 2020, 2021; Lee & Holyoak, 2023), suggesting that deliberation might sometimes cause an apparent preference reversal if the agent considers new information that it initially neglected. In standard accumulation-to-bound models, deliberation starts off with total ambivalence about preference, precluding the very possibility of a change of mind / preference reversal.

One way to potentially resolve this issue and enable this class of model to provide a measure of SoA would be to allow each option to enter deliberation with its own specific prior value estimate and precision. Mathematically, this would entail initializing the evidence variable at a point of non-neutrality. Previous work has presented sequential sampling models with such a starting point bias, representing prior beliefs about which option is *a priori* more likely to be the better one, based on instruction (Mulder et al., 2012), choice history (Urai et al., 2019), category preference (Lopez-Persem et al., 2016), personality trait (F. Chen & Krajbich, 2018), or choice history during embodied decisions (Kane et al., 2023; Lepora & Pezzulo, 2015; Molano-Mazón et al., 2024; Priorelli et al., 2024). A starting point bias could also be used to capture option-specific priors (Lee, Daunizeau, et al., 2023). These priors would form immediately upon presentation of the choice options, during the time it takes for perceptual processing to inform the decision apparatus about the identities of the options, but before explicit and intentional value deliberation begins. This would align with research showing that value signals in the brain arise automatically (Lebreton et al., 2009), even when not relevant (e.g., during perception). The formation of priors in this way would not only explain the starting point bias, but also the non-decision time (NDT), which is sometimes included in sequential sampling models to represent the time required for stimulus encoding or visual pre-processing (Fontanesi et al., 2019; Mulder & Maanen, 2013; Nunez et al., 2017). Once the priors were established, the evidence accumulation process at the core of the model would begin, terminating when the desired evidence level was reached. The difference in the posterior value estimates of the options (i.e., those at the end of deliberation) could then be compared to the difference in the priors to calculate a measure of SoA. In this way, this type of model could also account for changes of mind, because the chosen option would not always be the option with the higher prior value estimate.^2^

The mechanisms presented above would result in positive SoA on average (Lee & Daunizeau, 2021), and they would also account for the robust relationship between choice ease (absolute difference in value estimates, adjusted by value certainty) and SoA observed in the empirical data. But if deliberation were formalized as a process of recognition (or purification) of pre-existing value representations (Ratcliff, 1978), the signals that inform the initial and final preferences would have identical means – it would only be the precision that would differ. So, any SoA that might arise would be purely due to noise. Previous studies have shown that mere statistical noise could indeed yield a positive correlation between choice difficulty and SoA (Alós-Ferrer & Shi, 2015; K. M. Chen & Risen, 2010; Izuma & Murayama, 2013). However, an analysis of empirical data from multiple previous studies supports a cognitive source of SoA rather than a purely artifactual one (Lee & Pezzulo, 2023).

A more satisfactory way to account for the SoA results would be to focus on the crucial distinction between easy decisions (during which choices largely reflect prior value estimates and hence there is little or no SoA) and difficult decisions (during which novel sources of evidence are considered and hence there is often greater SoA). Choice difficulty might then be defined not based on some comprehensive measure of value but rather on an incomplete measure of value based on whatever (presumably non-exhaustive) information was considered during the initial isolated rating task. Under this account, the intra-decision dynamics of evidence accumulation (e.g., the evolution of the momentary drift rate) would differ as a function of choice difficulty. During easy decisions, when the choice is between options with very different values, evidence based only on the priors might already surpass the response threshold, in which case the choice could be made immediately without the need for deliberation, or with little additional evidence (see (Alós-Ferrer, 2018; Caplin & Martin, 2016; Diederich & Trueblood, 2018; Pezzulo et al., 2013) for alternative models that allow for this sort of automatic choice)^3^. This would imply no significant differences between the initial and final preferences and hence no SoA for such easy decisions. Difficult decisions, in contrast, are purported to elicit higher levels of information processing (Lee & Daunizeau, 2020, 2021; Pezzulo et al., 2013), which would potentially perturb the means of the option value representations if newly-considered information is not fully congruent with previously-considered information. It could be the case that early information systematically differs from late information. For example, it has been shown that neural activity (i.e., information processing in the brain) related to more salient or important attributes begins earlier than that related to less salient or important attributes (Lim et al., 2018; Maier et al., 2020; Sullivan et al., 2015). Furthermore, it has been proposed that the information considered in the formation of prior values and during deliberation is distinct and develops by means of passive retrieval from memory or active sampling via mental simulation, respectively (Pezzulo et al., 2013). Because the information considered at the start and at the end of a decision will not necessarily be the same (and thus the standard assumption of sampling from stationary inputs is removed), the value estimate for each option (not only its precision) could change over the course of deliberation. This suggests that the evidence accumulation rate in sequential sampling models should perhaps not be taken as static, but rather dynamic across deliberation time. This motivates the use of an extension to standard models in the form of an intra-trial time-varying drift rate. This approach has previously been suggested using drift rates that alternate or change in discrete stages within each trial (Diederich, 1997, 2003; Diederich & Oswald, 2014; Krajbich et al., 2010) or continuously over time (P. L. Smith, 2000). A dynamic drift rate combined with the notion that deliberation tends to last longer for difficult versus easy decisions could directly account for the observation that SoA positively correlates with choice difficulty.

The notion of an evidence accumulation rate that is dynamic across deliberation time is not new. For example, it has been proposed that this rate should vary as a function of attention, biased in favor of the option that is visually fixated at any point in time (*attentional Drift-Diffusion Model* or aDDM; (Krajbich et al., 2010; Sepulveda et al., 2020; Yang & Krajbich, 2022)). Here, the drift rate oscillates between relatively high and relatively low, as gaze (and presumably, attention) shifts between the choice options. Other work has proposed a drift rate that changes mid-deliberation, as an additional attribute (related to both options) enters consideration (*start time Drift-Diffusion Model* or stDDM; (Maier et al., 2020; Sullivan et al., 2015)). Here, the drift rate changes at one specific point in time, as two attributes start to be compared rather than just one. Clearly, this could be expanded to include additional attributes, and thus additional changes in the intra-trial drift rate. This would partially align with *Decision Field Theory* (DFT) models, where the drift rate fluctuates across deliberation time as (latent) attention shifts from one attribute to another (Busemeyer & Townsend, 1993; Diederich, 1997). Here, however, the momentary evidence accumulation rate is a function of only one attribute at a time, rather than a set of simultaneously-considered attributes that expands over time (as in the stDDM). Regardless of the differences between these models, they all demonstrate ways in which the preferences between options that are constructed during deliberation might differ depending on how attention (across options or across attributes) is dynamically allocated. Therefore, it is possible, in theory, that these models could account for the SoA phenomenon.

Here, we show that by combining key insights from three models that already include dynamic evidence accumulation (aDDM, DFT, stDDM), sequential sampling models can be made capable of accounting for SoA. Our approach follows that of *nested incremental modeling* used in other fields of mathematical modeling (Perry et al., 2007). From the previous models, our proposed model borrows the following notions: from the aDDM, evidence weights for specific options that diminish (but do not disappear entirely) when attention shifts away from those options; from DFT, attention that shifts between attributes (not only between options); and from the stDDM, additional attributes that come into consideration as deliberation time advances. In addition, we extend the concept of attributes beyond the inherent features of the options themselves to include contextual aspects of the decision that might alter the mapping from attributes to value. For example, when shopping for a new car, one might consider the size attribute. Initially, it could be that larger size is always deemed more valuable, as the most salient aspects of size are most easily recalled (e.g., more spacious seats, more room for luggage). However, upon further reflection (or mental simulation (Pezzulo et al., 2013)), larger size might come to be valued less, as other aspects are recalled (e.g., less room in the garage, more difficulty in maneuvering). Such contextual aspects might not be considered if the initial information is powerful enough to result in a fast choice.

In summary, our proposed model would allow for momentary attention to focus on specific attributes of specific options, which would temporarily diminish information processing for all other attributes and all other options, with more (and more refined) attributes entering the competition for attention across deliberation time. In what follows, we present a computational formulation for such a model and describe the key predictions that it makes with respect to choice behavior.

## Model specification and theoretical predictions

The core foundation of our model is a standard evidence accumulation process, with:

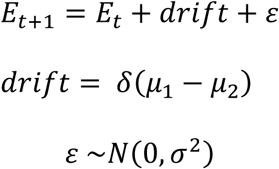

where *E* is the momentary evidence for one option over the other, *t* is an index for deliberation time (arbitrary units), 𝛿 is an agent-specific scalar representing evidence efficiency, 𝜇_𝑖_ is the value estimate of option *i*, and 𝜎^2^ is the strength of the white processing noise ε common to every decision.

Expanding this so that the process is based on multiple attributes is straightforward, replacing the overall value estimates 𝜇_𝑖_ with estimates of individual attributes 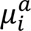 and the drift scalar 𝛿 with a vector of attribute-specific drift scalars 𝛿^𝑎^:

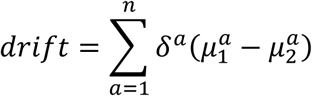

where *n* is the dimensionality of the attribute space for the decision at hand. Note that the scaling vector 𝛿^𝑎^ is mathematically equivalent (and notationally simpler) to a vector of attribute weights multiplied by a general scaling factor specific to the agent and common across attributes.

The next component of our model allows for the relative weights of the attribute dimensions to vary across time, as the attentional allocation of the agent fluctuates. The drift rate will thus change across time:

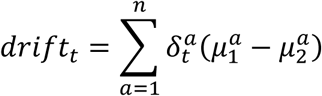

with 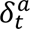 equal to 𝛿^𝑎^ for all momentarily active attributes and zero otherwise.

All that remains to complete the computational description of our model is to specify how the attribute weights 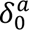 evolve across time. We will start by setting 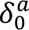 = 0 ∀ 𝑎 ∈ 𝐴 = {1,2, …, 𝑛}. For clarity, we will assume that only one attribute dimension will become active at a time (i.e., 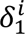 > 0 *and* 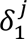 = 0 ∀ 𝑗 ∈ 𝐴\i). The dimension that becomes active first will correspond to the attribute that is generally the most salient for a particular choice context. For example, when choosing a snack food to consume, the tastiness attribute typically activates before other attributes like healthiness (Lim et al., 2018; Maier et al., 2020; Sullivan et al., 2015). An attribute might also activate sooner if it is more salient for a particular set of choice options, such as the attribute that has the highest individual assessment for any of the options within the set (Lee, D’Alessandro, et al., 2023), or because it is retrieved from memory during the recall of previous decisions or episodic *future thinking* (Schacter et al., 2007). The magnitude of an attribute weight on the choice (i.e., the contribution of evidence regarding that attribute towards the eventual choice) will be partially determined by the amount of attention the agent allocates towards that dimension. Here, the attention need not be conscious in the sense that the agent could report it if queried, but rather in the more general sense of the focus of information processing. Attention will be automatically attracted towards more inherently salient dimensions, and it can be adjusted towards dimensions that are more important to the specific agent or decision context.

Whether additional attributes (beyond the initial, most salient ones) will become active, and the point in time at which this would occur, will depend upon how informative^4^ the attributes already under consideration are with respect to the decision at hand (or in other words, the value of information or VOI that they offer; see (Behrens et al., 2007; Kobayashi et al., 2021; Pezzulo et al., 2013)). If the subset of currently attended attributes provides sufficient evidence that a specific option is the best, the agent might make its response without probing additional attribute dimensions. On the contrary, if the agent is unable to sufficiently distinguish the options based on the current active attribute subset, it might start to consider (i.e., allocate attention towards) an additional attribute – or proceed to simulate novel outcomes that may inform the decision. Here, we consider attention allocation to encompass both overt attention (e.g., shifting gaze between choice options or their attributes, as with the aDDM) and covert attention (e.g., thinking about option attributes based on memory). Furthermore, if the agent has already considered a particular attribute and concluded that that attribute dimension was unhelpful in distinguishing the option values, it might cease to consider that attribute for the remainder of the deliberation. This should be a function of the precision with which the agent has assessed the current attributes. For each estimate of the value of an attribute, an agent will also have a sense of precision about that estimate. Low precision will encourage further information processing, whereas high precision might discourage further information processing (as it would be redundant and thus inefficient to continue).

To better illustrate the process, take for example a simple choice between two snack food options: a chocolate bar and an orange. Let us assume that the most salient attribute of either option is its tastiness, so the agent will first consider that dimension. Imagine that after a brief deliberation, the agent realizes that both options are very similar in terms of tastiness and that it is difficult to choose which it prefers. The agent might then start to consider the healthiness of the options. If the agent were highly certain about its evaluation of the tastiness of the options, and thus highly certain that it would be of no use to consider that dimension further, it might replace tastiness with healthiness as the sole attribute under consideration (at that point in time). Otherwise, it might start to consider both attributes. Clearly, this process could iterate until either all relevant attributes had been considered or a subset of attributes had successfully provided enough evidence that either the chocolate bar or the orange was better than the other. Alternatively (or additionally), the agent might be motivated to mentally simulate the experience of selecting either option, potentially adding context-dependent attributes (e.g., the chocolate bar might melt if kept in the agent’s pocket, the orange might get moldy if not eaten soon) to the active set.

Below, we describe a novel computational model (which for brevity, we label the *salient attributes DDM* or *saDDM*) that formalizes the choice process described above. A key prediction of the model is that the spreading of alternatives (SoA) between choice options arises *during* a decision, as an effect of non-stationary integration of inputs from different attributes, which might vary during the decision process. Furthermore, the model predicts clear positive SoA for difficult choices, but lesser SoA for easy choices. The model also predicts that SoA would be greater for choices between options with higher attribute disparity, as the salient attributes would more likely differ between options and thus more attributes might be considered earlier in the deliberation process. Our simulations below will show that the saDDM accounts for empirical findings from previous studies that address SoA as a function of choice difficulty and attribute disparity (Lee & Hare, 2023; Lee & Holyoak, 2021). Apart from SoA, our model also correctly predicts the positive relationship that difficulty and disparity have with choice consistency, the negative relationship that difficulty and disparity have with response time, and the negative relationship that SoA has with response time (Lee & Hare, 2023; Lee & Holyoak, 2021). The model also outputs a relative contribution of each attribute to choice in line with the empirical data, even when the evidence efficiency scalars for each attribute are set to be identical.

## Methods

Using simulation, we demonstrate that the core predictions of our model qualitatively match all related empirical findings previously reported in the literature. We first generated choice trials based on experimental data pooled together across participants (n = 400) from two previous studies (Lee & Hare, 2023; Lee & Holyoak, 2021). Specifically, we used as input to our simulations the ratings that each participant provided for the first attribute (pleasure or *P*), the second attribute (nutrition or *N*), and the overall value (*V*) of each option, as well as the pairings of left and right options that each participant encountered on each choice trial. We calculated the difference in overall value (*dV*) for each choice trial as the difference between the option pair measures of overall value (option 1 minus option 2). We calculated the differences in the first and second attributes (*dP* and *dN*, respectively) in the same manner. Note that in the data table we worked with, we rearranged the data so that option 1 was always the one with the greater overall value rating. We used the estimated coefficients from a regression of overall value on the attributes (separately for each participant) as the participant-specific attribute weights. For each option on each trial, we calculated a measure of attribute disparity (orthogonal to *dV*) for each trial, as defined by Lee and Holyoak (Lee & Holyoak, 2021):

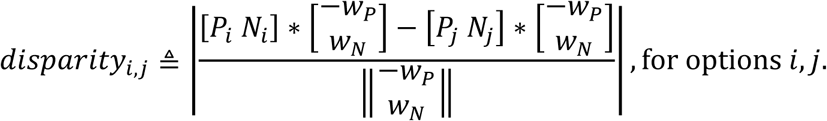

where *P* and *N* refer to the ratings with respect to the first and second attributes, respectively, and *w* refers to the weight of the contribution of each attribute to overall value.

Next, we simulated our evidence accumulation process for each trial for each participant. We set the response threshold magnitude to 1. For each participant, we set the drift rate scalar with a random draw from a normal distribution with mean 0.005 and standard deviation 0.001. For each participant, we set the standard deviation of the diffusion noise with a random draw from a normal distribution with mean 0.005 and standard deviation 0.0005. Following the norm in previous studies, we included *non-decision time*, which we set for each participant with a random draw from a normal distribution with mean 500 and standard deviation 100 time steps. For the start time of evidence accumulation for nutrition relative to pleasure (*relative start time*), we included a fixed amount and a variable amount that was proportional to one minus the difference in nutrition ratings for the options on each trial. Thus, relative start time was a function of the salience of nutrition for a given choice set. For each participant, we set the fixed and variable relative start time parameters with random draws from a normal distribution with mean 1000 and standard deviation 200 time steps. This set of parameter values yielded realistic output that was qualitatively similar to the experimental data, such as the RT distribution and the average choice consistency (defined as choices aligned with pre-choice value ratings) for difficult (low dV) and easy (high dV) choices. (The qualitative patterns remained robust across a wide variety of parameter combinations.) We set the maximum number of time steps to 10,000 to prevent excessively long simulation time. This is in line with analyses of experimental data of this type, where trials with response time greater than 10 seconds are often excluded. We deleted all trials where the simulation timed out, which left us with 13,414 trials (out of the original 13,820).

The evidence accumulation process on each trial proceeded as follows. Time and net evidence were initialized to zero. With each time step (*t*) beginning from trial onset, an increment of evidence equal to drift * dP was added to the net evidence while t was less than relative start time. Beginning at t = relative start time, the incremental evidence at each time step was equal to drift * (dP + dN). White Gaussian diffusion noise was added to the incremental evidence at each time step. The process stopped when the magnitude of the net evidence reached the response threshold. The choice (*Ch*) response on each trial was recorded as option 1 if the final net evidence (*E*) was positive or option 2 if the final net evidence was negative (coded as 1 for option 1, 0 for option 2). RT was recorded as the final time step (*T*) on each trial plus non-decision time, divided by 1,000 to make the quantities representative of seconds. Spreading of alternatives (SoA) on each trial was calculated as follows:

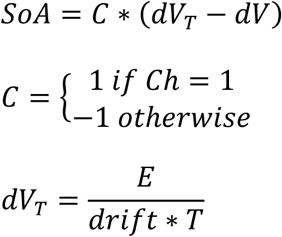

In this way, dVT can be thought of as the “inferred” value difference based on total accumulated evidence and deliberation time, whereas dV can be thought of as the “anticipated” value difference based on previous evaluations. The formula for SoA includes C because dV is calculated with respect to option 1 minus option 2, but SoA is defined with respect to the chosen option minus the rejected option.

Using the simulated data, we tested for relationships between the relevant decision variables: dV (as a measure of difficulty, with smaller values indicating greater difficulty), disparity, consistency, RT, and SoA. Specifically, we ran four separate regression analyses using the *fitlme* function in Matlab. First, we regressed consistency, log(RT), and SoA (separately) on dV and disparity. We then regressed log(RT) on dV and SoA. Finally, we regressed choice on dP and dN. We included random effects (both slopes and intercepts) for each individual experiment (7 in total) and for each individual participant (400 in total) in all regression models. For comparison with the results simulated from our model, we also ran the same set of regressions using the experimental data from previous studies (Lee & Hare, 2023; Lee & Holyoak, 2021).

## Results

As an initial check that our simulated data sufficiently resembled the experimental data, we examined the degree to which choices were consistent with reported option ratings (consistency averaged across participants). For difficult choices (defined as having dV less than the median dV for each participant), the average consistency was 67% in the experimental data and 66% in the simulated data. For easy choices (dV greater than the median), the average consistency was 93% in the experimental data and 90% in the simulated data. We also examined response times. For difficult choices, the average RT was 2.2s in the experimental data and 2.5s in the simulated data. For easy choices, the average RT was 1.8s in the experimental data and 1.4s in the simulated data.

Previous studies have reported several relationships between the different choice variables that we consider in this work. We first tested for a relationship between dV and disparity and consistency, RT, and SoA, since previous studies showed that both independent variables had a clear relationship with these dependent variables (Bhatia & Mullett, 2018; Lee & Hare, 2023; Lee & Holyoak, 2021). We thus regressed consistency, log(RT), and SoA (separately) on both dV and disparity. Our simulated data qualitatively reproduced all relevant patterns in the empirical data (see Figure 1).

**Figure 1:**
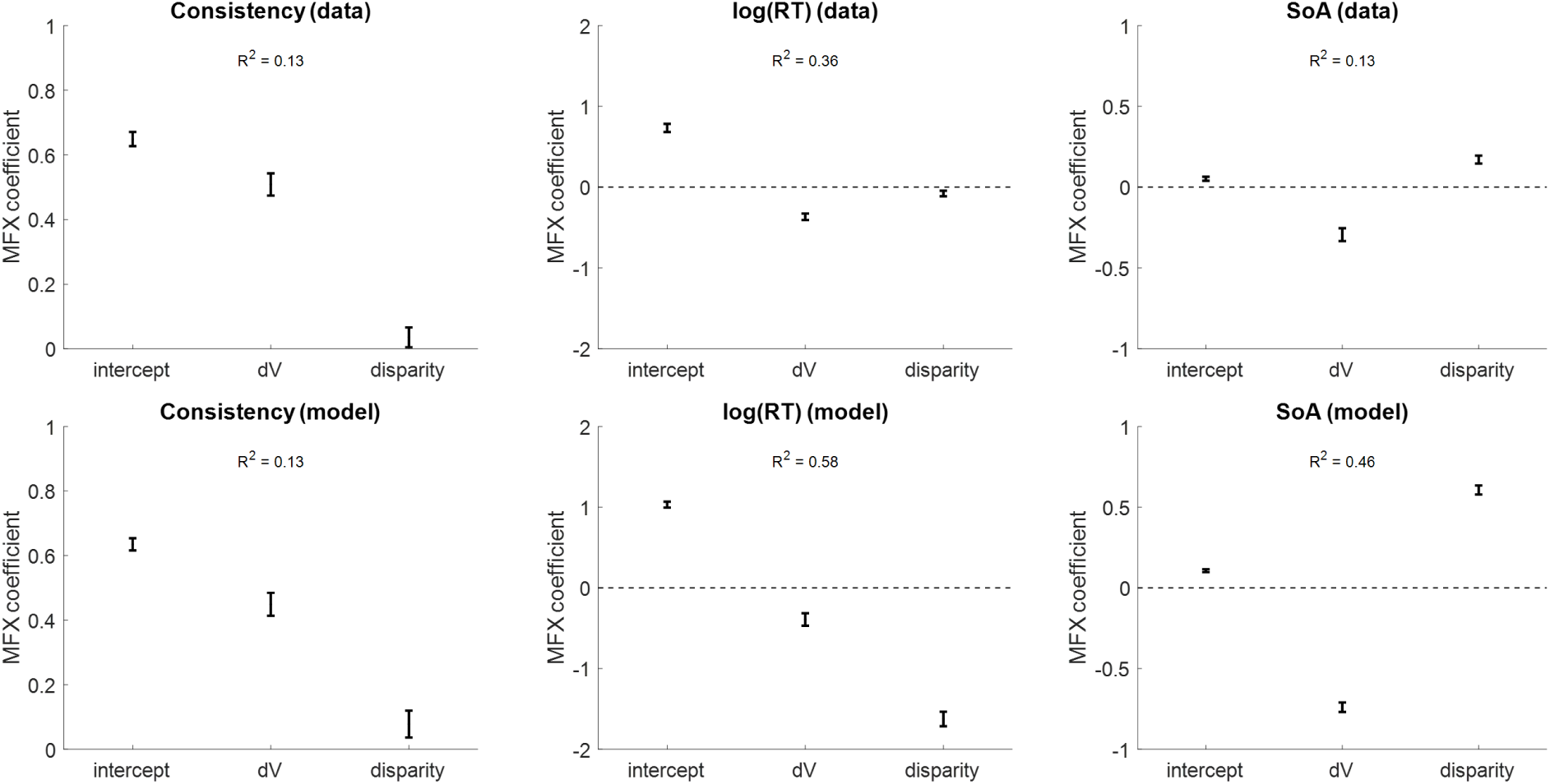
Relationships between dV, disparity, and dependent choice variables. The top row of plots shows the relationships that dV and disparity have with choice consistency, RT, and SoA in previously-reported experimental data. The bottom row of plots shows the same relationships in our simulated data. Error bars represent 95% confidence intervals.

We then regressed log(RT) on dV and SoA, since previous studies showed that SoA had a clear established relationship with RT that is qualitatively similar to the relationship between dV and RT (Lee & Hare, 2023; Lee & Holyoak, 2021, 2023; Lee & Pezzulo, 2023). Our simulated data qualitatively reproduced the pattern in the empirical data (see Figure 2).

**Figure 2:**
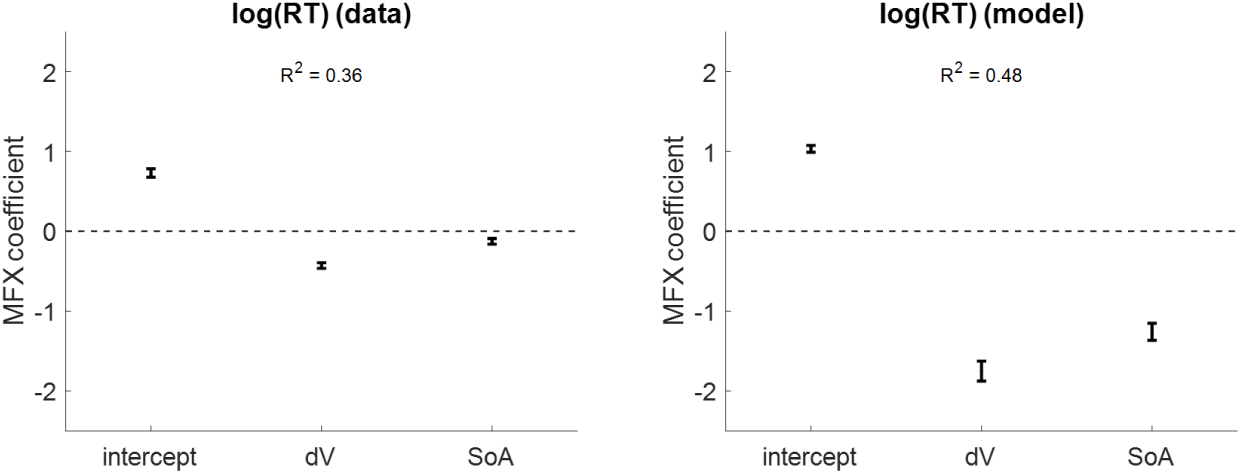
Relationships between difficulty, SoA, and RT. The left plot shows the relationships that dV and SoA have with RT in previously-reported experimental data. The right plot shows the same relationships in our simulated data. Error bars represent 95% confidence intervals.

Finally, we regressed choice consistency on the differences in the individual attributes on each trial (dP and dN). Our simulated data qualitatively reproduced the pattern in the empirical data (see Figure 3).

**Figure 3:**
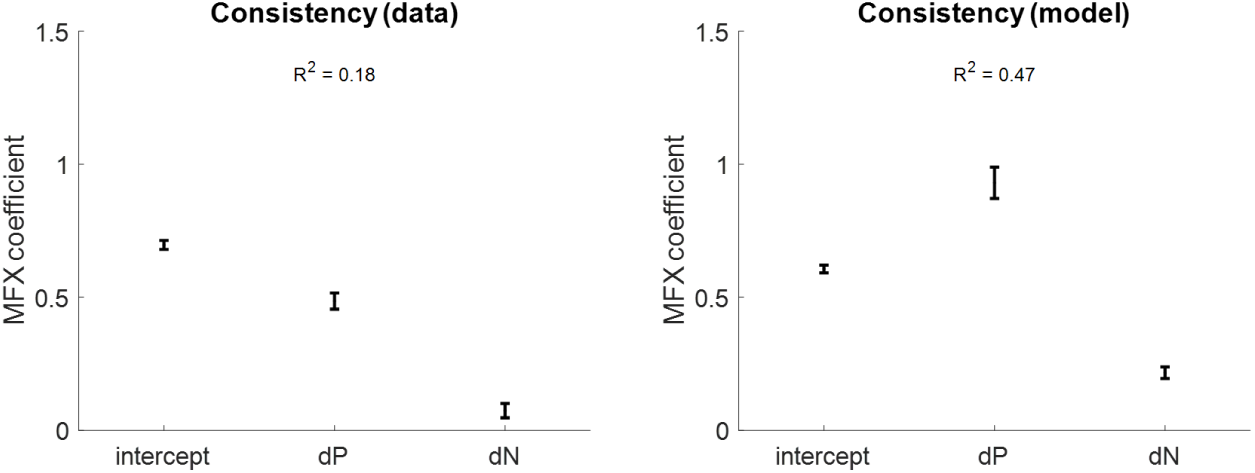
Relationships between dP, dN, and consistency. The left plot shows the relationships that dP and dN have with consistency in previously-reported experimental data. The right plot shows the same relationships in our simulated data. Error bars represent 95% confidence intervals.

Different aspects of the results we present above could arise under the previous models from which we drew inspiration: DFT, stDDM, aDDM. They are, after all, conceptually very similar models, depending on interpretation. DFT is based on accumulating evidence for different (single) attributes at different times across deliberation, when the agent shifts its focus. Under the stDDM, the agent fundamentally shifts its focus from one to two attributes at some point during deliberation. Under the aDDM, the agent could potentially shift its focus to different attributes (rather than to different options) at different points in time. So, in summary, the drift rate in each of these models would vary across time, and the output of the models could show some of the same SoA effects as we report for our saDDM. The key factor, regardless of which specific model formulation one chooses to use, is that the information about individual attributes that is incorporated into the final choice can vary across deliberation time. Another key factor, unique to our model, is that the degree to which different attributes are considered depends on the salience of those attributes with respect to the specific options under consideration in a particular decision.

## Discussion

In this work, we proposed a simple extension to classic evidence accumulation-to-bound models of value-based decision-making that can simultaneously account for response probability, response time (RT), and choice-induced preference change (as measured by the spreading of alternatives or SoA). We focused on the drift-diffusion model (DDM) because of its simplicity and analytical tractability, and because of its widespread success in accounting for standard decision variables (response probability and RT). Our computational model provides a principled account for how SoA could arise during choice deliberation by integrating insights from previous models of dynamic evidence accumulation (aDDM, DFT, stDDM), none of which explained SoA *per se*. Indeed, we are the first to directly explore such a possibility.

Our model (the *salient attributes DDM* or saDDM) assumes that the decision processes in the brain begin deliberation with imprecise representations of the values of each option, likely constructed from a subset of relevant attributes. Evidence starts to accumulate for only the most salient attributes at the beginning of deliberation, and evidence related to other attributes starts to accumulate at some later time point (Lim et al., 2018; Maier et al., 2020; Sullivan et al., 2015). We show via simulation that this is sufficient to give rise to SoA, complete with the correct (i.e., in line with the empirical findings) relationships between SoA and both choice difficulty and attribute disparity, as well as between RT and SoA. According to our theoretical model, the attention of the agent can switch between attributes over the course of deliberation (Busemeyer & Townsend, 1993; Diederich, 1997), and attention will modulate the rates of evidence accumulation (Krajbich et al., 2010; Yang & Krajbich, 2022). These theoretical elements of our model yield dynamic intra-trial evidence accumulation rates that could often result in posterior value estimates (e.g., at the time of choice response or during a post-choice rating task) that differ from the prior value estimates (e.g., at the beginning of choice deliberation or during a pre-choice rating task). Thus, the well-established SoA phenomenon can be fully explained by a principled process-based model, removing the need to rely on more descriptive accounts such as the post-choice resolution of cognitive dissonance (Brehm, 1956; Festinger, 1957).

The computational process we propose provides an account for how SoA between choice options might arise *during* a decision. However, such intra-decisional changes in evaluations are typically not observable (but see (Lee & Holyoak, 2023)). For this reason, experimental evidence for SoA generally comes in the form of isolated evaluations of the choice options, which are (on average) more distant after the choice task relative to before. Under the model described above, one would need to assume that pre-choice evaluations are made based on either a smaller subset of attributes than the subset that determines post-choice evaluations or a set of attributes that is more biased towards a particular subset. In the example above, this might mean that the chocolate bar and the orange were both initially (pre-choice) rated very highly (in isolation), based predominantly on their respective tastiness. During a decision in which these options were set against each other, however, the tastiness attribute might not have enabled a clear choice and so the healthiness attribute might have been considered more before the orange was chosen as the best option. Then, when the items were later re-evaluated in isolation (for the post-choice ratings), the orange might have been rated higher than the chocolate bar, because now the healthiness attribute (which favored the orange) was more influential in the overall ratings than it was before the choice. We focus here on identifiable attributes, but in a more general sense, the idea is that earlier assessments (either in isolation or in tandem) are less information-rich than later assessments, provided that the agent has proper motivation to seek enrichening information (Bénon et al., 2024; Lee & Daunizeau, 2020, 2021). Most likely, the posterior beliefs about option values from one task will inform the prior beliefs for a subsequent related task, following standard Bayesian computations.

One feature of the saDDM that sets it apart from other potential formulations is its ability to match the empirical relationship between attribute disparity and SoA. In brief, SoA is highest when the attribute composition of the options in a choice set is very different (high disparity), all else equal. To account for this, the saDDM relies on an assumption that disparity relates to the relative salience of the individual attributes across options. With low disparity, attribute composition is similar across options. Therefore, the most salient attribute is likely to be the same for all options. This would allow decisions to be more likely to be made based on only that mutually-salient attribute. On the contrary, with high disparity, attribute composition is dissimilar across options. Therefore, the most salient attribute is likely to be different for different options. This would make it more likely that decisions would be made based on multiple attributes, which would give rise to SoA (because pre-choice ratings would have been made based on only the attributes that were salient for each isolated option). In sum, the *basis of valuation* will differ as a function of disparity. SoA will occur primarily when the basis (mathematically, the set of attribute dimensions that comprise the value space where options are evaluated) changes between isolated ratings and comparison with alternative options, which is exactly the situation of high disparity.

When an agent initially considers each option in isolation (e.g., during a pre-choice rating task), different attributes might be more or less salient for different options. In that case, for some options, the value estimates at the start of the decision might relate more to one attribute, whereas for other options, the initial value estimates might relate more to a different attribute. Additionally, when choosing with a specific goal in mind, there could be a default primary attribute that the agent would consider before the others regardless of what the options were (Lim et al., 2018; Maier et al., 2020; Sullivan et al., 2015). Such an attribute should always be salient. These ideas combine to suggest that both option-specific factors and task-related factors could come into play to give rise to SoA. Predicting the change in attribute evaluations that might occur would be more complex, as some SoA could cause a particular attribute to have more impact on the choice while other SoA could cause the opposite. A future study dedicated to examining the rating process might help to better explain the intricacies of these various factors.

The saDDM relies on the assumption that post-choice ratings will resemble the latent value estimates that are constructed during choice deliberation up until the time a choice is made. This idea would generally align with the idea that value estimates are updated in a Bayesian manner during deliberation (Tajima et al., 2016), where the posterior estimates would endure without decaying back to the priors. It has been shown that the SoA effect is maximal *during* choice deliberation, although it then partially diminishes during a subsequent rating task (Lee & Holyoak, 2023). This suggests that post-choice ratings might be based on some combination of the information on which pre-choice ratings were based and the information on which choices were based. It seems that during isolated evaluations after choices have recently been made, the agent will tend to focus on the inherently salient attributes for each option but will also consider those attributes that contributed to the basis of comparison during the relevant choices. It is as if thinking about non-salient attributes during choice deliberation made those attributes more likely to be activated when considering the options again in the future (even in isolation). This would confer choices some history dependence that is not generally accounted for by current models. Future work should investigate the transient nature of SoA, as well as the neural underpinnings of bases of comparison during pre-choice rating, choice, and post-choice rating tasks.

Most studies related to SoA, including this one, consider only changes in ratings of overall value, but are agnostic about ratings of individual attributes. However, it has also been shown that SoA occurs even at the level of individual attributes (Lee & Holyoak, 2021). In other words, attribute evaluations tend to favor the chosen options more after the choices are made, relative to before (Lee & Holyoak, 2021). Perhaps SoA at the attribute level is caused by the same mechanisms that we have proposed for SoA at the level of overall value. This could be formalized as weights of sub-attributes (onto attributes and subsequently onto overall value) that change across deliberation time (or during choice versus rating tasks). Note that certain contextual aspects of value-based decision making could be related to higher-level attributes being broken down into sub-attributes in the mind of the agent (e.g., size could merely be a proxy attribute for the actual attributes involved in choosing a car, which might include comfort, utility, capacity, convenience, and ease of use). So, it could be said that deliberation involves not only the consideration of new attributes, but also the refinement of previous attributes (in our model, these could both be represented in the same mathematical format). Future work should explore sub-attributes and in what ways their relationship to higher-level attributes mirrors the relationship of attributes to overall value. This might build upon previous work showing that describing an attribute in different ways to highlight different objectives related to that attribute can alter preference judgments (Mertens et al., 2020; Ungemach et al., 2018).

The theory advanced in this paper contributes to the broader ongoing debate amongst researchers of value-based decision-making as to how the notion of value is represented in the minds of decision makers. The idea that value-related representations emerge at the time that they are called upon (e.g., when evaluating individual options or choosing between multiple options) is itself not widely contested. The debate is rather about *how* the value representations emerge. Some believe that option values are stored in memory and retrieved when needed (Padoa-Schioppa, 2011). While this hypothesis is intuitive, it is difficult to reconcile with the frequent observation that values reported for the same options differ depending on context. Others believe that value is constructed anew with each new task, built by an agglomeration of assessments of individual attributes of the option under consideration weighted by the importance of each attribute within the context where it is being evaluated (Bettman et al., 1998; Bhatia & Stewart, 2018; Johnson & Payne, 1985; Levav et al., 2010; Payne et al., 1988, 1993; Russo et al., 1996; Shah & Oppenheimer, 2008). Still others hold that memories of previous interactions with the option under consideration are retrieved (or imaginary interactions are simulated) and compiled into summary statistics (mean and variance across the sample of memories; (Ronayne & Brown, 2017; Stewart et al., 2006)). Another possibility is that value, at least in the standard sense used in neuroeconomics, might not be encoded in the brain at all (Hayden & Niv, 2021). More broadly, the pervasiveness of SoA could encourage skepticism that value or attribute estimates for choice options are encoded in the brain in the static form typically assumed by neuroeconomic models (Hayden & Niv, 2021).

While each of the above perspectives holds some merit, there is another aspect that is not directly addressed by this particular debate: value representations need not be identical across tasks (e.g., rating versus choice) – representations could have higher or lower fidelity depending on the task. In the ideal, there may be no reason for the value of an option (to a specific individual in a specific context) to be task dependent. However, cognitive operations are not cost-free (H. A. Simon, 1957; Zénon et al., 2019). Therefore, it is reasonable to believe that the sophistication of value representations formed with the purpose of enabling different tasks should align with motivational factors associated with each task. Specifically, the agent should conserve cognitive resources when establishing a value representation, investing only enough to satisfactorily perform the task at hand (Lee & Daunizeau, 2021; Sims, 2016; Warren et al., 2011). In other words, the agent might intentionally neglect some information in order to balance against the costs of information processing (Caplin & Dean, 2015; Fudenberg et al., 2018). What constitutes satisfactory performance is likely to differ between rating and choice tasks (for example). For an isolated individual rating, it might suffice simply to ascertain the valence of the option – the agent will want to approach the option if it has a value greater than zero, but to avoid the option if it has a value less than zero. Knowing that the rating will not be entirely isolated, in the sense that many different options need to be individually assessed, might induce a somewhat more refined value representation – the agent will want to accurately report its preferences, but will not be motivated to be highly precise. For a choice between options, on the contrary, the agent might not be able to perform the task satisfactorily (i.e., with a high enough feeling of confidence) unless it refines the value representations of the options by allocating additional cognitive resources (Lee & Daunizeau, 2021). Of course, this should only be the case if decisions are relatively difficult (i.e., when low-fidelity value representations would not be easily discriminable). In sum, we propose that value representations for individual ratings will be lower fidelity than for choices (in general). We also propose that value representations that emerge during choice deliberation will initially be lower fidelity (similar to that for ratings) but will convert to higher fidelity as necessary (i.e., if the decision is initially too difficult).

At this point, our theory suggesting that decision makers might use more- or less-refined value representations in a task-related way is speculative. However, the theory is not entirely novel if one considers an analogy within the field of perceptual learning, where it has been shown that greater discrimination difficulty induces more distinct representations of images (Ahissar & Hochstein, 1997). While the focus of this previous work is perception and not value, it shows an effect of task difficulty on the specificity of mental representations. This is very related to our line of thinking: for easy tasks, such as providing isolated ratings, imprecise representations could suffice; for difficult tasks, such as choosing between similarly-valued options, more precise representations may be needed. Future studies might address whether more- or less-refined representations as a function of difficulty do indeed also emerge in the brain during value-based decision-making.

Finally, while we have consistently referred to information as it relates to the choice options, it could also relate to the environment in which the agent finds itself. For example, the same option might seem highly valuable when examined on its own, yet only mediocre if the environment contains many superior options. Previous studies have exposed a so-called *distinction bias*, wherein an agent will be overly sensitive to variations in an attribute in terms of its contribution to the overall value estimates of the options (Hsee et al., 1999; Hsee & Zhang, 2004).

This results in ratings that differ when options are evaluated independently versus in tandem with other options, in line with the SoA phenomenon. It is as if the agent values both options equally if aware of only one of them at a time, but upon further reflection (encouraged by the need to choose between them), notices the differences between them and refines its value estimates accordingly.

## Code Availability

The analysis code used in preparation of this manuscript is publicly available on the Open Science Framework at https://osf.io/2bs73/.

## Footnotes

^1^This statement would no longer hold if the Tajima et al model were augmented with option-specific prior values (Tajima et al., 2016).

^2^One further assumption that this approach would require is that the process resulting in individual value estimates for each option (during the pre-choice rating task in a typical experimental paradigm) is similar to that which establishes the initial imprecise value representation at the start of a decision (because SoA is measured experimentally based on pre-choice ratings, as any latent value representation that forms at the beginning of a choice is not observable). Although this assumption might seem unreasonable, it may nevertheless be reasonable to assume that the process that yields isolated ratings lies somewhere between the very imprecise prior estimates and the relatively precise posterior estimates (i.e., it is a similar process but with an intermediate level of precision).

^3^Alternatively, in tasks where participants are required to report a decision by reaching or clicking with the computer mouse one of two response buttons located far away from the “start” position, the prior could determine the initial direction of (finger or mouse) movement, which may eventually be revised online during deliberation (Barca & Pezzulo, 2012, 2012; Dotan et al., 2018; Lee, D’Alessandro, et al., 2023; Lepora & Pezzulo, 2015).

^4^In this work, we do not distinguish between pure information, misinformation, or manipulated information. It is entirely possible that an attribute could be misinformative in the general sense (e.g., if someone believes that vitamins are bad for their health and thus are negatively influenced by nutrition). Similarly, the information content of an attribute could be manipulated by someone other than the agent (e.g., a used car salesman might persuade a shopper to focus more attention on a less important attribute, which might lead to a choice that would not maximize subjective value). Here, we refer to an attribute as *informative* simply if it informs the formation of a choice response.

